# Compact and contactless reflectance confocal microscope for neurosurgery

**DOI:** 10.1101/2020.05.13.093435

**Authors:** Jiahe Cui, Raphaël Turcotte, Karen M. Hampson, Matthew Wincott, Carla C. Schmidt, Nigel J. Emptage, Patra Charalampaki, Martin J. Booth

## Abstract

Visual guidance at the cellular level during neurosurgical procedures is essential for complete tumour resection. We present a compact reflectance confocal microscope with a 20 mm working distance that provided <1.2 µm spatial resolution over a 600 µm × 600 µm field of view in the near-infrared region. A physical footprint of 200 mm × 550 mm was achieved using only standard off-the-shelf components. Theoretical performance of the optical design was first evaluated via commercial Zemax software. Then three specimens from rodents: fixed brain, frozen calvaria and live hippocampal slices, were used to experimentally assess system capability and robustness. Results show great potential for the proposed system to be translated into use as a next generation label-free and contactless neurosurgical microscope.

## 1. Introduction

The primary aim in the treatment of brain cancer is maximal tumour resection with minimal damage to adjacent brain tissue. Numerous technologies have been developed to provide real-time intraoperative guidance in order to achieve this goal. In particular, specialised optical microscopes have shown to provide valuable morphological information during surgery [1–3]. However, most of these systems were not optimised to enable visualisation of individual cells. Low numerical aperture (NA) was used, normally below 0.07, to offer a working distance (WD) over 200 mm [4], and thus allow space for surgical manipulations.

To overcome the limitation in spatial resolution, endoscopic probes have emerged and proven effective in delineating the tumour margin with higher spatial precision [5–8]. In addition to improving the discrimination between diseased and normal cells, endoscopic probes are more compact than most specialised surgical microscopes, making them convenient for deployment in operating rooms. However, some practical complications have also been encountered during their use in clinical applications [2, 5]. One major drawback is that endoscopic devices require direct contact with tissue. They are therefore susceptible to erythrocyte contamination, and frequent cleaning of the probe tip is unavoidable [5]. Another caveat originates from the difficulty to stabilise and apply uniform pressure on the probe when held by hand [2].

In sum, it would be ideal to combine the highly desirable features of both endoscopes and specialised surgical microscopes into a single, compact system such that cellular level resolution can be achieved while eliminating the need for direct contact between instrument and tissue. This configuration would further allow for manipulation of surgical tools below the microscope objective using both hands as in conventional surgical microscopes.

Apart from system configuration, the employed contrast mechanism also affects the feasibility of adopting an imaging method for intraoperative use. Fluorescence guided surgery, as one of the most commonly exploited techniques, has provided useful information on the cellular and molecular scale for brain tumour resection [2, 9–11]. However, the use of fluorescent dyes in the human brain is limited by many factors, such as tumour selectivity, fluorophore uptake due to the blood-brain-barrier, fluorophore brightness and toxicity, photobleaching, injection timings, dose constraints [12–14]. Consequently, only three markers have been approved for neurosurgical procedures in humans [13]. Therefore, novel label-free intraoperative devices are in high demand.

Reflectance confocal microscopy (RCM) has for long been of special clinical interest due to its favourable optical sectioning abilities and label-free nature [15–18]. The implementation of a pinhole in both the illumination and detection paths helps to eliminate out-of-focus light adjacent to the focal plane. Without being constrained by the excitation spectrum of a specific fluorophore, the choice of illumination wavelength is primarily dependent on optical properties of the tissue itself, with additional considerations for spatial resolution. Furthermore, compared to fluorescence microscopy, the signal level for RCM is usually much higher for a given incident power. Less light is required at the tissue, reducing potential possibilities of photodamage; while stringent requirements imposed on the detection sensitivity of fluorescence imaging can be loosened as well [11]. Most importantly, owing to a contrast mechanism based on differential refractive indices, RCM has recently been found to provide information on cellularity, architecture, and morphological features simultaneously for tumour discrimination in *ex vivo* studies [18], showing great potential for surgical translation.

In this paper, we present a novel compact reflectance confocal microscope with a 20 mm WD that operates in the near-infrared and provides <1.2 µm spatial resolution over a 600 µm × 600 µm field of view (FOV). We first introduce our design concepts before presenting the microscope setup in detail. Then, the performance of the optical design is evaluated using commercial Zemax software. System capability and robustness is systematically assessed in three biological specimens. First, 60-µm thick fixed mouse brain is used to validate spatial resolution and imaging fidelity in biological tissue. Optical sectioning abilities are then demonstrated through axial-stack imaging and three-dimensional (3D) reconstruction of frozen mouse calvaria. Live tissue imaging capability is verified through maximum intensity projections of 100-µm deep axial-stacks in the rat organotypic hippocampus. Finally, novelties as well as space for future development of the system is discussed in view of the practical needs for next generation neurosurgical microscopes.

## 2. Materials and methods

### 2.1. Design concept

The design of our confocal reflectance microscope aims to overcome limitations of currently available neurosurgical systems, while considering the ease of translation from bench-top to clinic. First, a system WD sufficient for preventing direct contact with tissue and permitting insertion of surgical tools was a main priority. Another major goal was to provide a spatial resolution of ∼1 µm across the whole FOV such that cellular level examination could be accomplished. Furthermore, system compactness was given special consideration to enable easy manoeuvring of the microscope in the operating theatre. Finally, to make this system affordable and reproducible, off-the-shelf rather than custom-designed lenses were used.

### 2.2. System setup

The main microscope design is illustrated in Fig. 1(a). For reduced scattering and minimal blood absorption, a near-infrared (NIR) wavelength of 830 nm was chosen [19, 20]. A low coherence superluminescent diode (SLD, SLD830S-A20, Thorlabs) was used to suppress speckle noise without increasing pinhole size [21]. Light was coupled into the system via a single mode fibre and power after the objective was set to 5 mW for all experiments. To minimise detection of backscattered light from components along the optical path, a quarter-wave plate (QWP) was implemented in combination with a linear polariser (LP) and a polarising beam-splitter (PBS). The LP generated horizontally-polarised illumination light, which transmitted through the PBS, before passing the QWP whose fast-axis was oriented 45^°^ relative to the polarisation direction. The PBS then reflected vertically-polarised signal light into the detection path while any spurious horizontally-polarised light would be eliminated. A compact gimbal-less dual-axis MEMS scan mirror (S31155, Mirrorcle Technologies) with a clear aperture of 2 mm was integrated for system miniaturisation, providing a maximum mechanical scan angle (MSA) of ± 4.8^°^ in both the horizontal and vertical directions. The mirror was conjugated using four lenses (L1, 30120, effective focal length (EFL) = 25.4 mm, Edmund Optics; L2, 49363, EFL = 175 mm, Edmund Optics; L3, 49361, EFL = 125 mm, Edmund Optics; L4, 49360, EFL = 100 mm, Edmund Optics) to the back aperture of a 20× magnification, 0.42 NA, 20 mm WD, air immersion plan-apochromat objective (LWDO, MY20X-804, Mitutoyo). In the detection path, backscattered signal light was coupled into an adjustable aspheric fibre collimator (FC2, CFC-11X-B, Thorlabs) with a focal length of 10 mm. A single mode fibre with a core size of 5 µm, equivalent to 0.7 Airy unit (AU), was used as a confocal pinhole to couple signal light into a silicon avalanche photodetector (APD, APD430A/M, Thorlabs). Components within the main microscope body were separately mounted on a 200 mm × 250 mm breadboard (MB2025/M, Thorlabs) and further enclosed in a protective casing. L4 and the QWP were mounted into lens tubes that extended from the microscope body and could be easily unscrewed for portable transit. A fibre-in fibre-out (FIFO) configuration was implemented for miniaturisation. The system footprint was kept to 200 mm × 550 mm and the physical layout of the system with a magnified view indicating the insertion of a scalpel between the microscope and sample is depicted in Fig. 1(b).

**Fig. 1.**
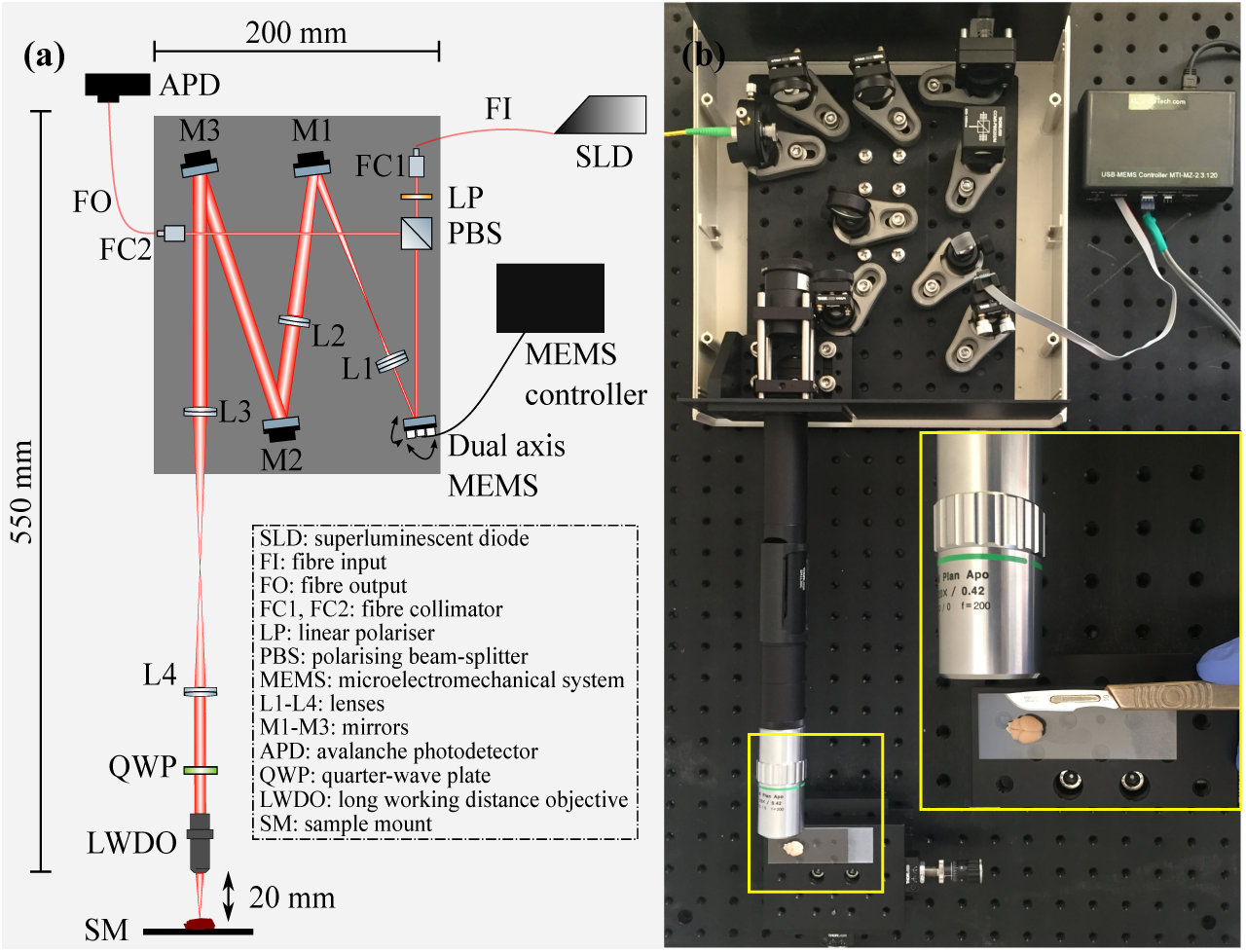
Microscope design and physical layout. (a) Schematic drawing of the imaging system. Components are individually labelled and full description is given in the dotted box. (b) Physical layout of microscope system taking up a 200 mm × 550 mm bench-top footprint. The inset shows a magnified view indicating the insertion of a scalpel between the microscope and sample.

Scanning control and image acquisition were performed using custom-developed Python software via a standard laptop (XPS 15 7590, Dell) and FPGA (USB-7855R, National Instruments). Analog triangle wave scan signals were sent to the MEMS mirror controller and the analog detection signal from the APD was used for image construction. Distortion due to the non-linear velocity of the scan mirror was corrected using a pre-calibrated look-up table. Images were displayed on a custom GUI interface and later processed using ImageJ software.

### 2.3. Zemax evaluation of system aberrations

A compact optical system exclusively using commercial lenses is prone to aberrations, especially when large scan angles are required such that oblique beams enter short focal length lenses at peripheral positions. In order to determine a lens combination that causes minimal accumulation of system-induced aberrations, we modelled and evaluated the design using commercial Zemax software 18.9. As the objective lens model was not available, imaging performance was analysed at the objective back focal plane in afocal mode.

Ideally, diffraction-limited performance (DLP) is highly desirable in microscopic systems. However, depending on the application, this is not always necessary. For our purpose, cellular level resolution is easily achievable within a 1 µm regime. Substantial beam expansions were also required to fill the objective back aperture. Therefore, rather than aiming for DLP that was likely to be achieved within only a narrow region about the optical axis, we slightly relaxed the criterion to obtain more uniform image quality across the whole FOV. A Strehl ratio of > 0.7 was sustained for an equivalent FOV of approximately 600 µm × 600 µm. Assuming a linear relationship between MSA and FOV using small angle approximation, system performance was evaluated at a series of MSAs across the full scan range with increments equivalent to 100 µm at the sample. The Strehl ratio, diffraction encircled energy, and Huygens point spread function (PSF) were individually assessed.

#### 2.3.1. Strehl ratio

The Strehl ratio (SR) is commonly used as a performance criterion for high resolution imaging systems. The theoretical SR variations with different MSAs across both the *x*- and *y*-directions are shown in Fig. 2(a), where the *x*- and *y*-axis are horizontal and vertical to the base plate, respectively. The Maréchal criterion of SR = 0.8 is used as reference for DLP. It can be seen that at all scan angles along both the *x*- and *y*-axis, the SR is kept well within 0.7. A symmetrical variation can be noticed across the FOV in both scan directions, most likely to have originated from the balancing of field flatness distortion at different lateral positions when optimising for the highest uniform SR. Comparison between the *x*-scan and *y*-scan curves indicate similar degrees of system-induced aberration such that the same SR is achieved for a certain scan angle.

**Fig. 2.**
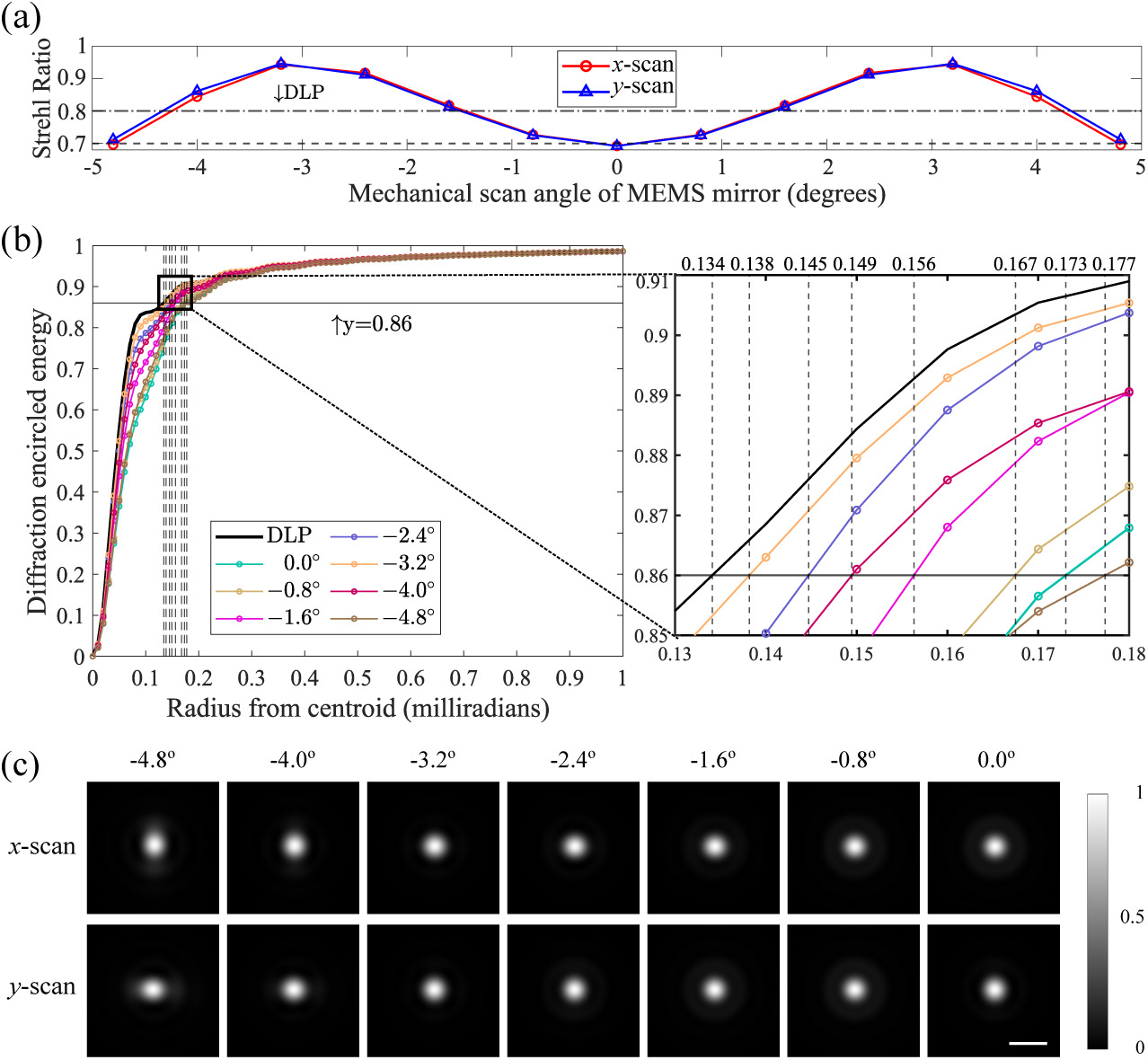
Zemax evaluation of system imaging performance. (a) Theoretical SR variation as a function of MEMS mirror MSAs in both *x*- (red) and *y*-directions (blue). DLP of 0.8 SR at the objective back aperture and the system optimisation result of 0.7 SR for all MSAs along both scan directions are indicated with horizontal lines. (b) Diffraction encircled energy versus radius from centroid in milliradians for different MSAs. Black curve is shown for DLP. Solid black line marks 86% of the total energy. Magnified plot highlights radius values at which the diffraction encircled energy reach 86% of the total energy for DLP and all other MSAs. (c) Huygens PSF profiles across a half FOV for both *x*- and *y*-scans. Intensity is individually normalised. Scale bar: 200 milliradians.

#### 2.3.2. Diffraction encircled energy

Variations in the illumination PSF’s size and energy distribution is examined by plotting the diffraction encircled energy diagram, as demonstrated in Fig. 2(b). Each curve refers to the gain of cumulative energy with radius for a different MSA, and that for DLP is highlighted in black as reference. Only results for negative scan angles are plotted due to the symmetry witnessed in Fig. 2(a). It can be found that all scan angles follow a similar steep accumulation trend of energy comparable to that of DLP. To assess the degree of deterioration in PSF size brought by relaxation of the SR, we calculate the PSF’s full width at half maximum (FWHM) at different MSAs by first taking the 1/e^2^ width of the PSF, with in which 86% of total beam power is contained, and further considering a conversion factor of 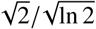. The radius values in milliradians at which the diffraction encircled energy curves reach 86% of the total energy for DLP and all other MSAs are highlighted in the magnified plot of Fig. 2(b). Considering a focal length of *f* = 100 mm for L4, and a 20× demagnification introduced by the objective lens, as well as assuming a similar intensity PSF (IPSF) for both the illumination and detection paths in a confocal system, the effective system IPSF 1/ e^2^ width and FWHM are calculated and given in Table 1.

**Table 1.**
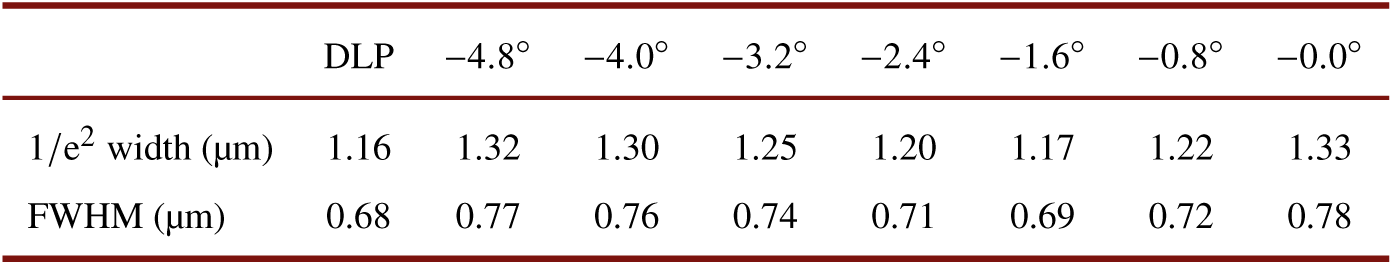
Theoretical 1 / e^2^ width and FWHM of effective IPSF for DLP and different MSAs in the proposed confocal system.

Results show that despite system optimisation resulting in an SR of 0.7, the illumination IPSF FWHM across the whole 600 µm × 600 µm FOV is successfully kept within a ∼1 µm regime, and the largest deviation relative to DLP is ∼250 nm, a negligible amount for cellular level imaging.

#### 2.3.3. Huygens PSF

Finally, to understand the main aberration components in the system, the Huygens PSF, which is the diffraction-limited PSF using direct integration of the Huygens wavelets method, was assessed for half of the FOV along both *x*- and *y*-scan directions. It can be observed in Fig. 2(c) that for both cases, the shape of the PSF starts to deviate from an ideal circular spot for larger angles between −3.2^°^ and −4.0^°^ and becomes elongated in one direction, showing moderate astigmatism. Defocus and spherical aberrations are the main on-axis aberration components, as can be seen from the annular structures surrounding the bright central region. However, no significant variations in PSF profile are noticed in either directions, indicating a reasonably uniform image quality for the entire 600 µm × 600 µm FOV.

### 2.4. Reflectance imaging

Spatial resolution was calibrated using a stage micrometer with 1 µm line width (AGL4202, Agar Scientific) placed at the central FOV. Profile plot of the line showed a FWHM of 1.10 µm. Effective image intensity in a reflectance confocal microscope is given by I_eff_ = | (h_ill_ · h_det_) ⊗ o_r_ |^2^, where h_ill_ and h_det_ denote the illumination and detection amplitude PSFs (APSF) and o_r_ represents the object reflectance profile [22]. For simplicity, we use a Gaussian approximation for h_ill_, h_det_ and o_r_, and assume similar distribution of the illumination and detection APSF such that h_ill_ = h_det_ = h_a_. Further considering the relation of h_eff_ = |h_a_|^4^ between effective system IPSF h_eff_ and single path APSF h_a_ in a confocal system, the FWHM of h_eff_ was calculated to be 0.84 µm. We compare this experimentally derived effective IPSF with the result in Table 1 for FWHM at 0.0^°^ MSA. It can be seen that the built confocal system provided an effective IPSF consistent with that of the theoretical system, only ∼7% larger in size. Additionally, a spatial resolution of <1.2 µm could be resolved in the reflectance images.

During experiments, a manual translation stage (PT3A/M, Thorlabs) was used for 3D translation of specimens. Imaging took place with the microscope mounted both horizontally and vertically on a mechanical arm. No visible difference in image quality was noticed, indicating the microscope’s ability to perform reliably in all orientations.

### 2.5. Sample preparation

System capability and robustness were systematically validated using three biological specimens: 1) 60-µm thick fixed brain slice from a Thy1-GFP mouse; 2) excised mouse calvaria; 3) 350-µm thick organotypic hippocampal slice from a male Wistar rat. All animal work was carried out in accordance with the Animals (Scientific Procedures) Act, 1986 (UK).

Thy1-GFP mice were overdosed with pentobarbital before transcardial perfusion with phosphate buffered saline and 4% paraformaldehyde (PFA). Subsequently, the fixed brain was extracted and immersed in PFA for 24 hours. Finally, the fixed brain was cut into 60-µm thick coronal slices and mounted between a microscope slide and a 170-µm coverslip for imaging.

Calvaria was extracted from frozen mouse cadavers using scissors. The left portion of the calvarial bone between the frontal and coronal suture was then cut and glued onto a microscope slide for imaging. Imaging was performed in air.

Organotypic hippocampal slices (350-µm) were prepared from male Wistar rats (P7) and cultured on Millicell CM membranes (Millipore). Slices were maintained in culture media (Life Technologies/Sigma Aldrich), at 37°C for 7-14 days prior to use. Culture media was replaced every 2-3 days. Cultured slices were transferred to a petri dish, fixed with vacuum grease and kept in physiological Tyrode’s solution for imaging (in mM: 120 NaCl, 2.5 KCl, 30 glucose, 2 CaCl_2_, 1 MgCl_2_, and 20 HEPES; Sigma Aldrich; pH = 7.2-7.4; *n* = 1.33).

## 3. Experimental results

### 3.1. Ex vivo imaging of fixed mouse brain

Taking advantage of the well-known anatomy and morphology of the mouse brain, fixed slices were imaged to characterise system spatial resolution and imaging fidelity in biological tissue.

The longitudinal fissure, separating the left and right hemispheres, was located at the top-most border of the cerebral cortex (Fig. 3(a)). Figure 3(b) shows prominent dark neurons within the upper cortical region. Adjacent regions above and below featured much sparser distributions of scattered cells. The corpus callosum (CC) was recognised as a clearly discernable fibre tract in the lower cortical region (Fig. 3(c)), separating the outer cortex from the hippocampus. Just before reaching the CC, sparsely distributed neurons of varying sizes were witnessed. Figure 3(d) shows the CA1 field of the hippocampus. As expected, a neatly defined hypo-reflective cell body layer (CBL) was formed by CA1 pyramidal cells; while adjacent neuronal processes (NPs) were distinguished as distinct dark grooves projecting from the CA1 neurons in a parallel configuration, as well as regions with high reflective signal.

**Fig. 3.**
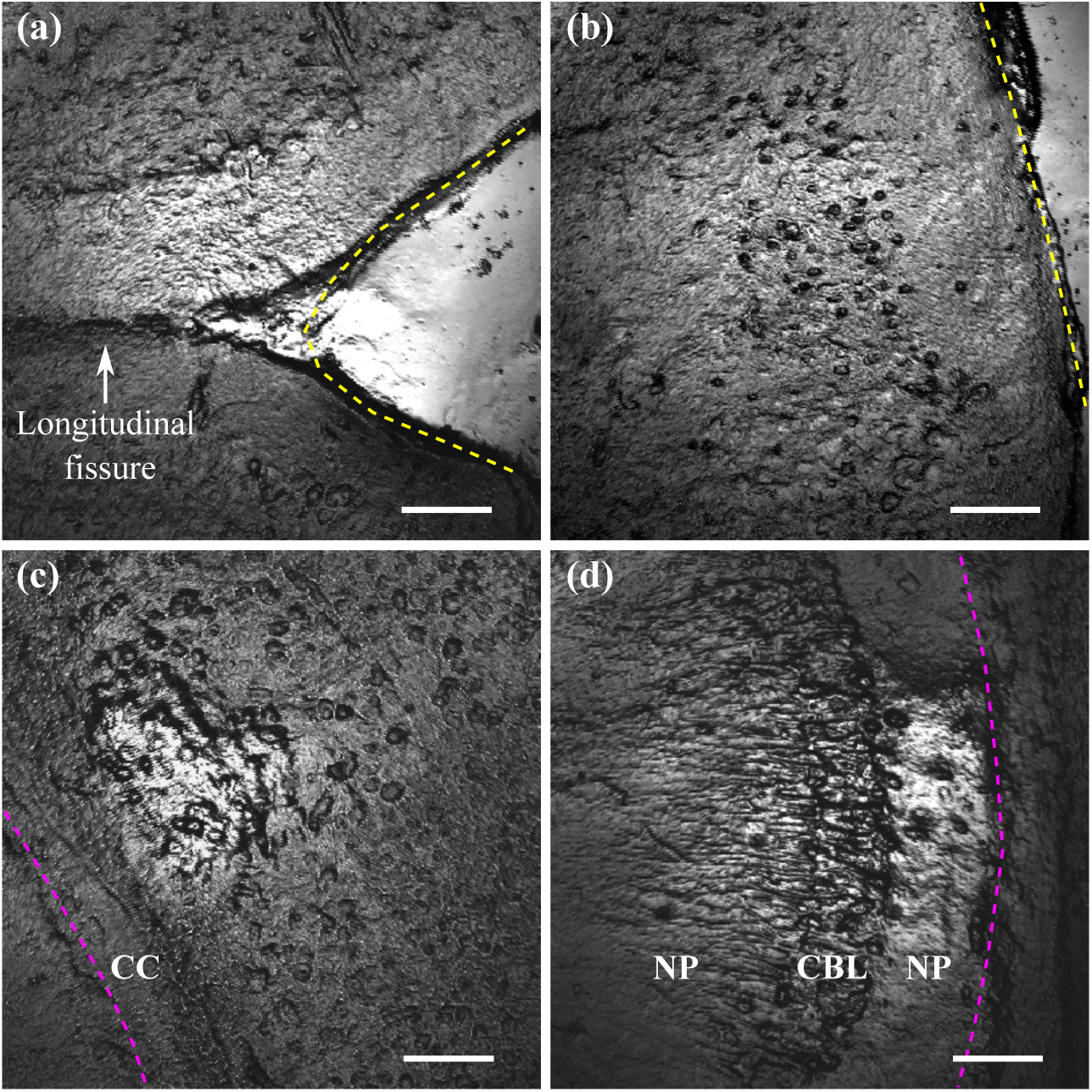
Imaging of 60-µm thick fixed brain slice from a Thy1-GFP mouse. (a) Top-most border of cerebral cortex showing longitudinal fissure. Yellow line: edge of sample. (b) Upper cortical region displaying densely packed dark neurons. Adjacent regions above and below featured sparser distributions of scattered cells. (c) Lower cortical region featuring the corpus callosum (CC) and neurons of varying sizes. Magenta line: outer border of the hippocampus. (d) CA1 field of the hippocampus. Pyramidal cells formed a neat hypo-reflective cell body layer (CBL). Neuronal processes (NP) were distinguished as distinct dark grooves projecting from the CA1 neurons in a parallel configuration, as well as regions with high reflective signal. Scale bar: 100 µm.

The results above suggested high spatial resolution and imaging fidelity of the microscope within biological tissue. The 170-µm coverslip is liable to introduce spherical aberration when used with an air immersion objective without coverslip correction. Although this effect is more detrimental with high NA objectives, especially in the presence of sample tilt [23], slight blurring effects are still likely to be present during imaging. Nevertheless, we have demonstrated that the proposed system utilising RCM, has the potential of imaging morphological features of the mouse cortex and hippocampus without any form of labelling [24, 25].

### 3.2. Ex vivo imaging of frozen mouse calvaria

The mouse calvaria is known to be highly scattering, and its significant surface curvature causes further deterioration of the light field for optical microscopy [26]. Here we show that our RCM system maintained sufficient optical sectioning, even when severe optical distortions were present, to achieve 3D imaging at the cellular level. Two different locations were imaged: the first where the bone surface was approximately normal to the optical axis, and the second where it was tilted by several degrees. Distinct features in hard bone and bone marrow demonstrated the axial sectioning ability when volumetric data was acquired with 5-µm axial steps (Fig. 4).

**Fig. 4.**
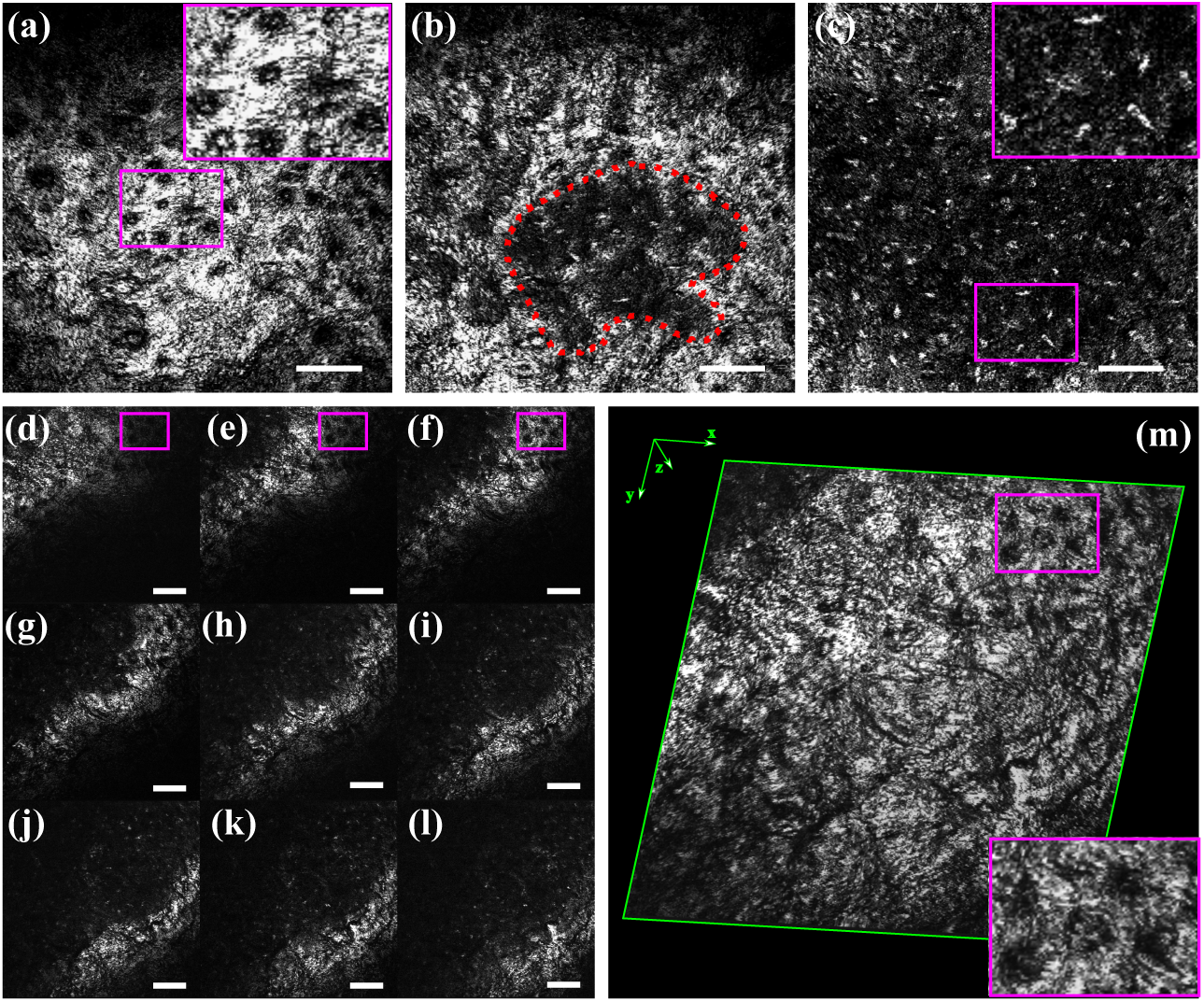
Volumetric imaging of the mouse calvaria at locations where the bone surface was approximately (a-c) normal to the optical axis and (d-m) tilted by several degrees. (a) In the hard bone, the locations of osteocytes were revealed by negative contrast from the bright hard bone matrix. (b) The top surface of a bone marrow cavity was observed 45 µm below the hard bone surface. A sharp interface was visible between the hard bone and bone marrow (dotted line). (c) A further 5 µm deeper, the bone marrow cavity was larger, occupying almost the entirety of the FOV. (d-l) Consecutive slices within the calvaria showing a distinct cross-section of the hard bone matrix at each depth. Osteocytes within magenta boxes gradually come into focus from (d) to (f). (m) 3D reconstruction of (d-l) showed a smooth transition of the bone surface. The similar osteocyte pattern as in (d-f) was observed. Scale bar: 100 µm.

At the first location, the hard bone matrix covered the entirety of the FOV (Fig. 4(a)). A magnified view of osteocytes is given in the inset, visualised by negative contrast with the strongly reflective collagen bone matrix [27]. The transition between hard bone and bone marrow was clearly discernable as a sharp and curved interface (Fig. 4(b); dotted line), confirming the microscope’s excellent optical sectioning ability, and suggesting an axial resolution of <5 µm within highly scattering biological tissue. The bone marrow cavity appeared to be less reflective than hard bone, likely as a consequence of the freezing process (Fig. 4(c)), while scattered bright cells were nonetheless clearly identifiable. Nine 5-µm thick stack images were taken before reaching the transition zone, indicating a hard bone thickness of approximately 45 µm.

At the second location, nine consecutive axial-stack images were taken (Fig. 4(d-l)) and due to surface tilt, only a distinct cross-section of the hard bone matrix was visible within the focal plane at each incrementing depth, showing a diagonal progression pattern from the top left corner to bottom right. After performing 3D reconstruction of the consecutive stack images, the surface of the hard bone matrix could be visualised (Fig. 4(m)), showing smooth transition and featuring the same osteocyte pattern indicated within boxes in Fig. 4(d-f). Note how the osteocytes gradually come into focus from Fig. 4(d) to Fig. 4(f).

The above results provided validation of the penetration capabilities of the microscope through optically dense and highly scattering media, as well as its optical sectioning capability across the whole FOV.

### 3.3. Live tissue imaging of the rat hippocampus

In the final experiment, to verify the live tissue imaging capability of the microscope, we chose a 350-µm thick organotypic hippocampal slice and left a thick layer of Tyrode’s solution between air and tissue to mimic an aqueous physiological interface.

Figure 5(a) illustrates the rat hippocampus highlighting three biologically significant regions, namely the dentate gyrus (DG), CA3 field, and CA1 field. A 100-µm deep axial-stack with 5-µm intervals was taken for each region and a maximum intensity projection (MIP) was performed to build the 100-µm thick volume. Demonstration of the maximum projected hippocampal slice is given with rotational views at 0°, 30°, and 60^°^ relative to the normal projection (Figs. 5(b) – (d)).

We started at the DG (Fig. 5(b)) which is acutely bent in shape. A dark granular cell layer was visible in the central region of interest. Adjacent regions of NPs showed high reflectivity and dense fibrillarity, forming strong contrast against the neuronal cell bodies. MIP of the CA3 field revealed a narrower layer of NPs towards the edge of the hippocampus, surrounding a region of dark pyramidal cells (Fig. 5(c)). Visible reflective signal contrast was also seen between the cell bodies and NPs in the CA1 field, though lower than its CA3 counterpart (Fig. 5(d)).

**Fig. 5.**
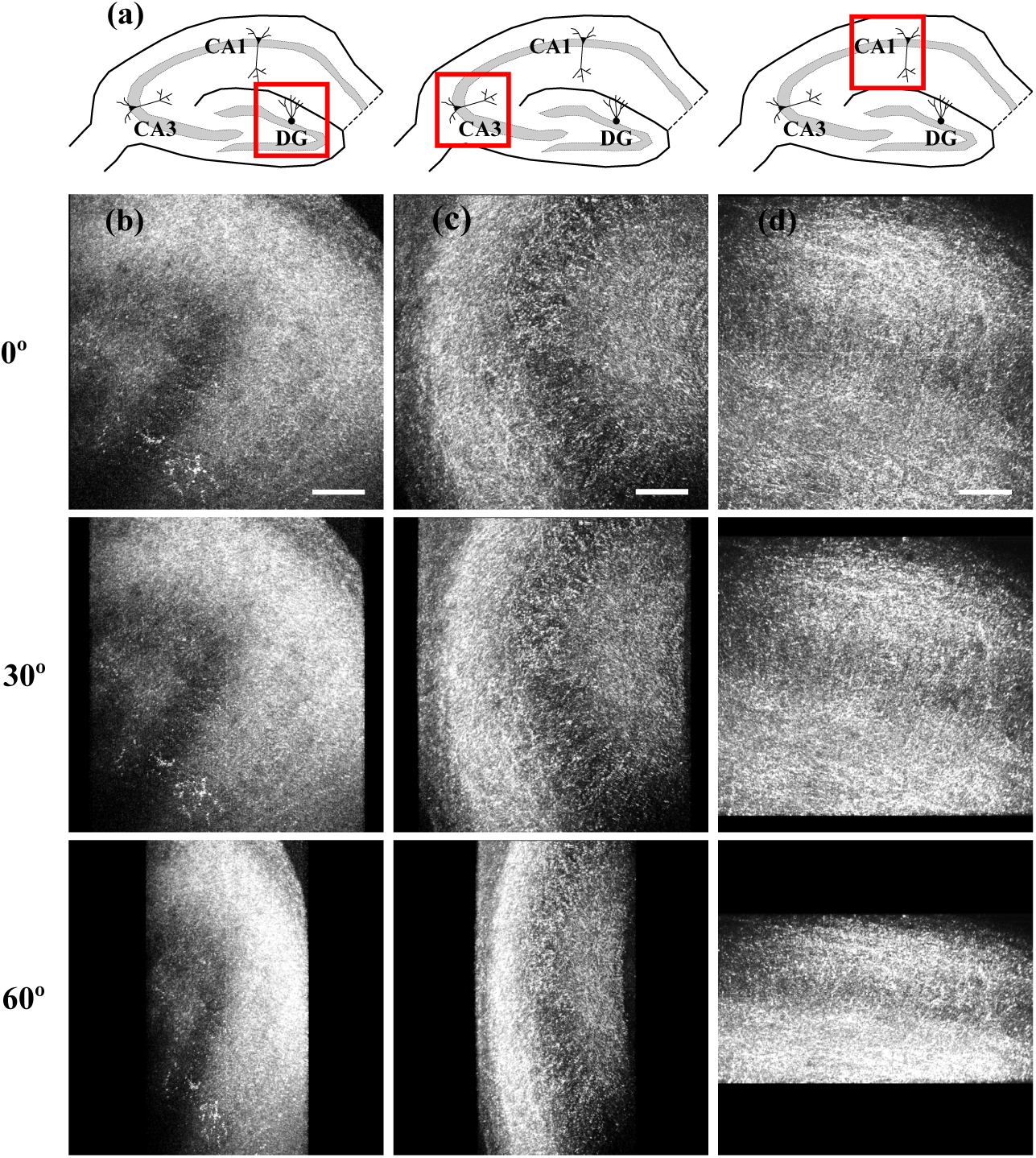
Rotational views of maximum projected 100-µm deep organotypic hippocampal slices at 0°, 30°, and 60^°^ relative to the normal projection. (a) Schematic diagram of the rat hippocampus. Solid red boxes indicate imaging regions for (b) – (d). DG: dentate gyrus. (b) MIP of the DG showed a dark granular cell layer and adjacent regions of NPs with high reflectivity and dense fibrillarity. (c) MIP of the CA3 field showed a narrower layer of NPs surrounding a region of dark pyramidal cells. (d) Visible reflective signal contrast was also seen between the cell bodies and NPs in the CA1 field. Scale bar: 100 µm.

Successful imaging of the live hippocampus up to 100 µm in depth validated the capability of the proposed microscope for live tissue imaging before translation to pre-clinical experiments. It should be mentioned that although RCM has been previously reported for live tissue and *in vivo* imaging of the mouse brain [28, 29], as well as imaging of the fresh human cadaver [30], rather than using a high NA objective lens for deep brain imaging, we have adopted a low NA (0.42) air immersion objective. In addition, compared with results obtained in Section 3.1, it can be found that due to the fixation method for *ex vivo* imaging, as well as a loss of aqueous tissue contents, different characteristics were observed between fixed and live brain tissue. This mainly took the form of a highly reflective background with individual dark cells seen in the fixed brain slice, as opposed to the dark background, bright fibrous filaments, and dark band of cells in the live hippocampus.

## 4. Discussion

Many factors influence the effectiveness of a device for image-guided brain surgery, which can be briefly split into two categories. The first incorporates intrinsic functionality features of the instrument, such as resolution, field of view (FOV), imaging depth, frame rate, contrast mechanism, etc. The second, however, is equally important but much easier to neglect, which is the fact that instrumentation should be designed in such a way that its full functional advantage could be exerted with minimum effort in real life scenarios.

The reflectance confocal microscope presented in this work fills a substantial gap in surgical practice. Specifically, key features of conventional surgical microscopes and endoscopic probes were combined such that cellular level resolution was achieved, while enough space was left between device and tissue to prevent direct contact and frequent probe cleaning, as well as allow for insertion of surgical tools. A minimal number of off-the-shelf components was used with considerations for compactness, affordability and reproducibility. Sufficient resolution was provided both laterally and axially to discern individual dendrites as well as delineate the border of tissue layers, and imaging was achieved at least 100 µm within live tissue, suggesting the system’s competence for clinical translation. Comparison between images of fixed and live brain tissue revealed instructive information on how variations in cell and tissue morphology could possibly change the refractive index to give different textures under the proposed microscope. This has interesting clinical implications and agrees with findings described in [15, 17, 18].

In addition, the concept entails some practical advantages. A combination of NIR illumination with fibre-based detection reduces the system’s susceptibility to room light interference, which is normally in the visible light spectrum [31], implying that surgical procedures could be performed under normal lighting conditions [11]. Longer wavelengths would further decrease interference of ambient light as well as increase the imaging depth [19, 29], but at the expense of a reduced spatial resolution. In addition, benefiting from a label-free and contactless concept, imaging requires no additional labelling of the tissue or excessive sterilization processes of endoscopic probes, enabling simplification of pre-surgical procedures.

Further system development should mainly address the following aspects: imaging speed and depth. First of all, the current frame rate is limited by the analog sampling rate to 1 Hz. The implementation of a scanning module with position feedback would enable operation in resonant mode and thus substantially increase the frame rate to around 6 Hz [32]. To achieve this, a high-speed digitiser could be used to increase capture speed, through which images of higher diagnostic quality could be displayed with pixel resolutions higher than 512 × 512. In the current system, for the maximum imaging FOV of 600 µm ×600 µm, resolution is limited by pixel count rather than optical restrictions. A faster scan mirror could further enhance imaging speed, but current integrated MEMS technology would require using mirrors with smaller clear apertures, thus needing larger beam expansions, ultimately introducing stronger system aberrations.

Second, the implementation of a wavefront shaping device, such as a deformable mirror (DM), would be extremely beneficial for performing simultaneous adaptive optics (AO) and remote focusing [33, 34]. Correction of both sample- and system-induced aberrations would allow a tighter focal spot to arrive at the specimen and more signal to be delivered through the detection pinhole so as to improve signal-to-noise ratio and permit deeper imaging into the highly heterogeneous brain tissue [35, 36]. The combination of a deeper imaging depth and a more accurate refocusing unit together with the optical sectioning abilities of the confocal microscope would be especially useful for finding the three-dimensional tumour margin.

## 5. Conclusion

We have developed a compact microscope using off-the-shelf components for guidance during neurosurgery. The system achieved a spatial resolution of <1.2 µm while leaving a 20 mm working distance between the device and tissue to prevent direct contact and enable easy manipulation of surgical tools. Taking advantage of reflectance confocal microscopy, the proposed system is label-free and possesses three-dimensional imaging capabilities. The potential for pre-clinical translation is experimentally validated in distinct biological specimens: fixed mouse brain, frozen mouse calvaria, and organotypic hippocampal slice. This study provides initial ground for future development towards next generation neurosurgical microscopes while also leaving space for prospects in other areas of surgical imaging.

## Funding

European Research Council (ERC) under the European Union’s Horizon 2020 Research and Innovation Programme (812998 and 695140).

## Acknowledgments

The datasets generated during and/or analysed during the current study are available from the corresponding author on reasonable request.

## Disclosures

The authors declare no conflicts of interest.

## Notes

### Competing Interest Statement

The authors have declared no competing interest.

